# GPR39 Localization in Aging Human Brain and Correlation of Expression and Polymorphism with Vascular Cognitive Impairment

**DOI:** 10.1101/2021.07.15.452525

**Authors:** Catherine M Davis, Thierno M Bah, Wenri H Zhang, Jonathan W Nelson, Kirsti Golgotiu, Xiao Nie, Farah N Alkayed, Jennifer M Young, Randy L Woltjer, Lisa C Silbert, Marjorie R Grafe, Nabil J Alkayed

## Abstract

**INTRODUCTION:** The pathogenesis of vascular cognitive impairment (VCI) is not fully understood. GPR39, an orphan G-protein coupled receptor, is implicated in neurological disorders but its role in VCI is unknown.

**METHODS:** We performed GPR39 immunohistochemical analysis in postmortem brain samples from mild cognitive impairment (MCI) and control subjects. DNA was analyzed for GPR39 SNPs, and correlated with white matter hyperintensity (WMH) burden on premortem MRI.

**RESULTS:** GPR39 is expressed in aged human dorsolateral prefrontal cortex, localized to microglia and peri-capillary cells resembling pericytes. GPR39-capillary colocalization, and density of GPR39-expressing microglia was increased in aged brains compared to young. SNP distribution was equivalent between groups; however, homozygous SNP carriers were present only in the MCI group, and had higher WMH volume than WT or heterozygous SNP carriers.

**DISCUSSION:** GPR39 may play a role in aging-related VCI, and may serve as a therapeutic target and biomarker for the risk of developing VCI.

## Background

Aging is the strongest risk factor for Alzheimer’s disease (AD) and related dementia [1]. With people increasingly surviving into their 80s, 90s and beyond, the number of people living with dementia is projected to triple by 2050[1]. Historically, vascular cognitive impairment (VCI), the second most common cause of dementia after AD, and AD were considered distinct entities[2]. There is a growing appreciation, however, of the overlap and similarities between AD-and VCI-related dementias[3]. Indeed, many patients with AD have vascular pathology[3], and brains from VCI patients also exhibit Alzheimer’s pathology. Furthermore, both entities often occur in the same individual and exacerbate each other. For example, AD patients with ischemic lesions have poorer cognitive function than those with no vascular lesions [4], highlighting the importance of studying cerebrovascular mechanisms underlying VCI.

While the pathogenesis of VCI is not fully understood, small vessel disease (SVD) is a key underlying factor [5]. Ischemic lesions resulting from SVD appear on T2-weighted MRI scans as hyperintensities primarily localized in the cerebral white matter, termed white matter hyperintensities (WMH). In a prospective study, we have shown that acceleration of WMH burden is an early, presymptomatic pathological change predictive of conversion to the clinical diagnosis of mild cognitive impairment (MCI) [6]. Furthermore, we have also confirmed that autopsy brains from deceased patients with dementia and histopathological evidence of SVD had larger WMH volume on premortem MRI [7].

The current study was designed to investigate the involvement of the orphan G-protein coupled receptor GPR39 in neurovascular injury underlying VCI. GPR39 has been implicated in a variety of neurological disorders[8], but its involvement in VCI has not been investigated. GPR39 binds zinc, which plays an important role in neurotransmission, and is implicated in the pathology of AD[9]. Zinc is necessary for formation of the toxic forms of amyloid beta, which in turn attenuates zinc neuronal signaling [9, 10], identifying GPR39 as a potential novel therapeutic target in neurological disorders including AD. GPR39 has also been implicated in a range of pathological processes other than amyloid pathology [9], and has been proposed to play a role in other neurological conditions, including depression and seizures. GPR39 has also been implicated in vascular pathology, including vascular inflammation and calcification [11, 12]. The expression and cellular localization of GPR39 in human brain is not known, and its potential link to VCI has not been investigated. Therefore, we set out to characterize GPR39 expression in human brain and its potential association with VCI.

To that end, we identified postmortem brain samples collected from former participants of the Oregon Brain Aging Study (OBAS) with a clinical diagnosis of MCI. As control, we used brains from age-and sex-matched subjects with no cognitive deficit, as well as young adults. We conducted two studies. We first performed immunohistochemical analysis to localize GPR39 and determine its correlation with VCI. In the second study, we determined if known GPR39 single nucleotide polymorphisms (SNPs) correlate with WMH burden on premortem MRI.

## Methods

### Study subjects

Thirty-five brain and 78 DNA samples were obtained from the Layton Aging and Alzheimer’s Disease Research Center Brain Bank Repository at Oregon Health & Science University. Distribution of samples between the two cohorts and demographic data are presented in Table 1.

**Table 1.**
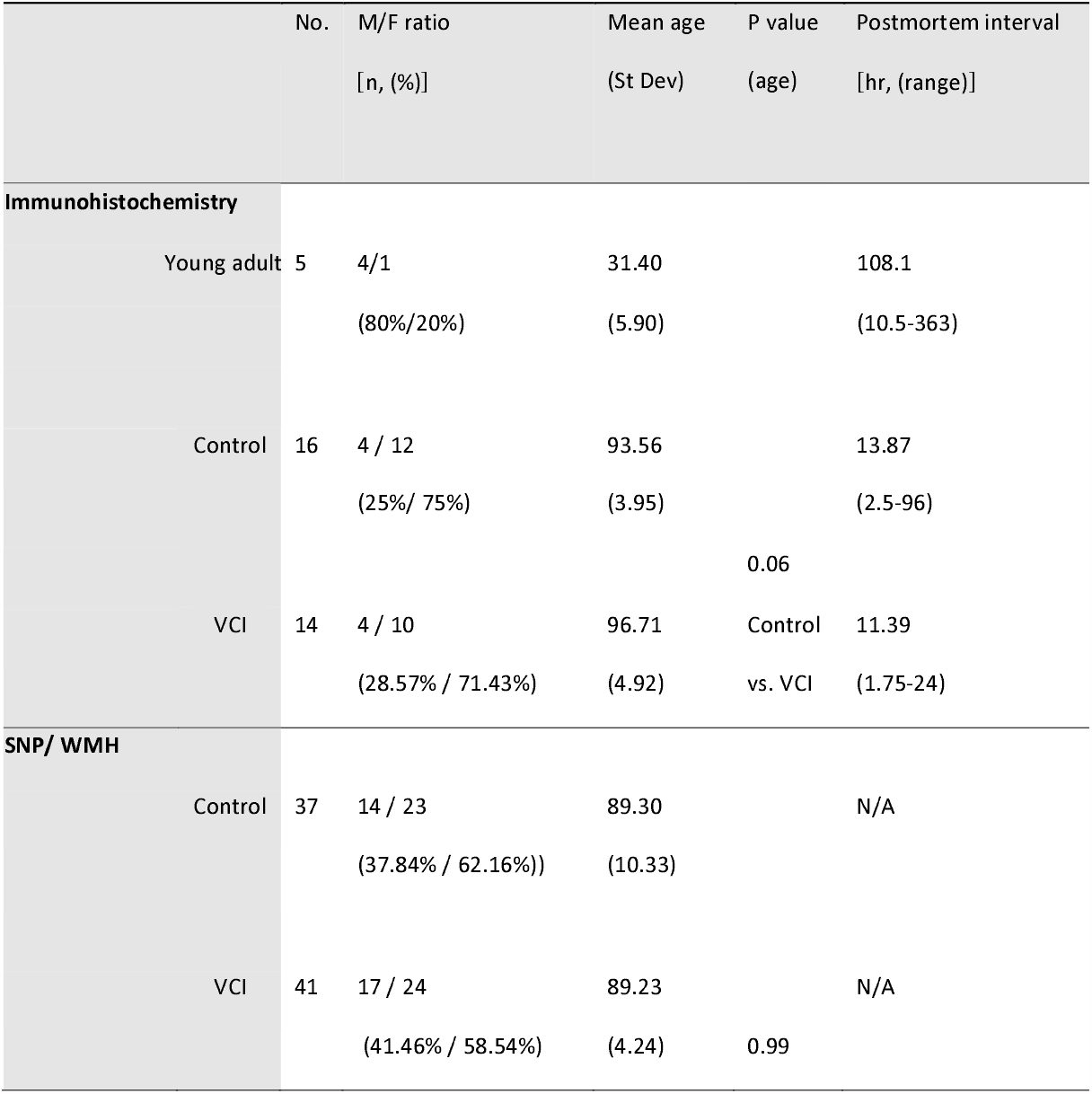
Demographic data of subjects for immunohistochemistry and SNP/WMH analysis.

For immunohistochemical assessment, postmortem tissues were selected based on a clinical diagnosis of MCI before death, or in the case of control subjects, on absence of cognitive impairment. The collection, storage and distribution of information and autopsy tissue were in accordance with guidelines established by the Layton Center Research Repository in compliance with Federal Regulations and Oregon Health & Science University policies.

For measurement of WMH volume and genetic analysis, subjects were selected from the OBAS [13] based on availability of premortem 3T MRI measurements, genomic DNA, and clinicopathologic diagnosis indicative of MCI. WMH volume measurements and clinical dementia ratings (CDR) were quantified as previously described[14].

### Immunohistochemistry

Formalin-fixed dorsolateral prefrontal cortex (dlPFC) sections were immunolabeled with antibodies against GPR39 (anti-GPR39, extracellular domain, antibodies online ABIN1048812), CD68 (anti-CD68, abcam ab955), collagen IV (anti-collagen IV, COL-94, ThermoFisher Scientific MA1-22148), NeuN (anti-NeuN, clone A60, Millipore Sigma MAB377), GFAP (anti-GFAP, clone2E8, abcam ab68428) and SMA (anti-actin, α -smooth muscle, Sigma A2547). Antigen retrieval was carried out by pretreatment with citrate buffer, pH 6 in a steamer for 30 minutes followed by 3% hydrogen peroxide to reduce endogenous peroxidase activity. For both CD68 and collagen IV ImmPRESS HRP anti-mouse IgG (peroxidase) polymer detection kit was used (Vector, MP-7402) and subsequently the reaction visualized using diaminobenzidine. For GPR39 ImmPRESS-AP anti-rabbit IgG (alkaline phosphatase) polymer detection kit was used (Vector, MP-5401) and subsequently the reaction was visualized by Vector blue alkaline phosphatase (Blue AP) substrate kit (Vector, SK-5300). For negative controls the primary antibody was replaced by species-appropriate serum.

### Image Acquisition and Analysis

Images were acquired on a Zeiss AxioScan.Z1 Slide Scanner. Resulting czi images were converted into tiff files after which grey and white tissue brain regions were delineated and separated into two different images using Adobe Photoshop; grey and white regions were separated as mutually exclusive regions and files. Images were broken down by tiling each image into an array of 8×8 tiff files (64/ image) using freeware Irfanview. Resultant files were processed using FIJI/ImageJ. Density of each marker was quantified by converting labeled area into a binary mask, within which particles were filtered (5-150 um in diameter), counted and divided by total tissue area. Co-localization was determined by quantifying overlapping regions of interests of each of the markers and expressed as a percentage of either CD68 or collagen IV. Image acquisition and analysis were carried out in a blinded manner.

### Genotyping

DNA was extracted from patients as previously described [7]. Two previously reported SNPs linked to vascular disease, rs13420028 and rs10188442 [15] were investigated. The genotype of each subject for SNP status (homozygous WT, heterozygous or homozygous for each SNP) for the gene that encodes for GPR39 was determined by allelic discrimination using TAQMAN technology (Invitrogen). SNP assays were purchased from ThermoFisher for rs10188442 and rs13420028. Quantitative PCR assays were performed on the QuantStudio Real-time PCR System (Life Technologies) and data collected using Applied Biosystems QuantStudio 12K Flex Software. DNA quality assessment and qPCR assays were performed in the OHSU Gene Profiling Shared Resource, according to manufacturer’s instructions.

### Statistics

For immunohistochemical data, one-way ANOVA with Tukey’s multiple comparisons test was used, with significance threshold set at p<0.05 using Prism 8 (GraphPad). Data are presented as mean +/-SEM. For SNP analysis, Fisher’s exact test was used, analysis was carried out using Stata Statistical Software, Release 16 (StataCorp LLC).

### Study Approval

Consent for research purposes specifically was obtained from next of kin at time of autopsy.

## Results

There were no significant differences in mean age and postmortem interval between the aged MCI and control subjects for either the immunohistochemical or the genotype and WMH analysis (Table 1).

### Immunohistochemical localization and expression of GPR39 in the young and aged human dlPFC

We first assessed whether GPR39 protein is expressed in the dorsolateral prefrontal cortex of the human brain, its cellular localization and distribution, and whether expression levels are altered in aging and in individuals with VCI. We found that GPR39 is robustly expressed throughout the dlPFC. We observe a non-significant trend for increased GPR39 in the aged dlPFC, with overall density of GPR39 immunoreactivity being similar in aged control and VCI, both in gray and white matter (Figure 1). In terms of distribution, we observed two populations of GPR39-positive cells, one in close proximity to blood vessels, and the other within the tissue parenchyma.

**Figure 1.**
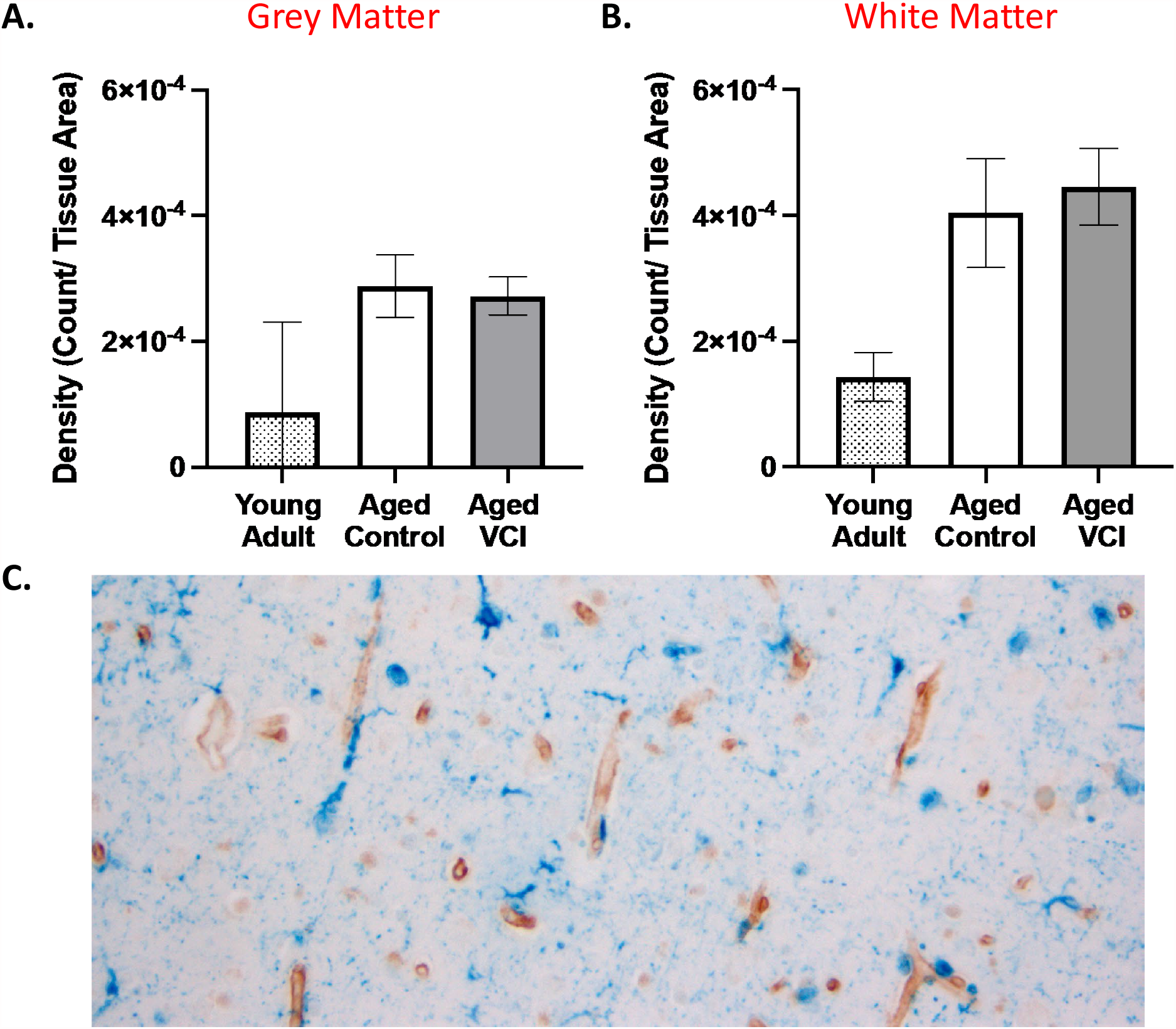
GPR39 expression in dlPFC of young and aged control and aged VCI human brains. Quantification of GPR39 expression density in both grey, **A**, and white matter, **B**, of young adult control and aged control and VCI human brains. **C**, Representative image of GPR39 immunolabeling (blue) of human aged dlPFC. Data represent mean ± SEM (n=5-15/group).

To investigate the cellular identity of GPR39-expressing cells in the human dlPFC, we performed double-labelling of brain sections for GPR39 and collagen IV, which labels the basal lamina of blood vessels (Figure 2). We observed that many, but not all of GPR39-positive cells are intimately associated with capillaries, with some cells colocalizing with capillaries (arrows), while others are in close proximity (peri-capillary; arrowheads). Based on morphology and close association with capillaries, this cell population is consistent with pericytes. We used multiple antibodies against two pericyte markers, human platelet-derived growth factor receptor beta (PDGFRβ) and human neuron-glial antigen 2 (NG2). However, none of these antibodies tested worked on our human formalin-fixed tissue. We were therefore unable to immunohistochemically confirm the identity of these GPR39-positive peri-capillary cells. These GPR39-positive cells were predominantly associated with capillaries (Figure 2C-D), and rarely seen in larger blood vessels (not shown), further supporting pericyte-localized GPR39 expression. Upon quantification, we observe a trend for decreased collagen IV density in the aged groups compared to the young adult control samples. Furthermore, we observe a striking increase in co-localization of GPR39 with collagen IV in the aged dlPFC, compared to young where colocalization is largely absent (Figures 2 A-B). In grey matter, co-localization is increased in both aged groups compared to young adult, increasing from 1 ± 0.02% co-localization in young adult to 32.21 ± 5.19% in aged control and 21.55 ± 3.14% in aged VCI (p<0.05, 1-way ANOVA with Tukey’s multiple comparisons test), with no significant difference between aged control and VCI. In white matter co-localization was increased in the aged control dlPFC compared to young adult (p<0.05, 1-way ANOVA with Tukey’s multiple comparisons test), again there was no significant difference between aged control and aged VCI.

**Figure 2.**
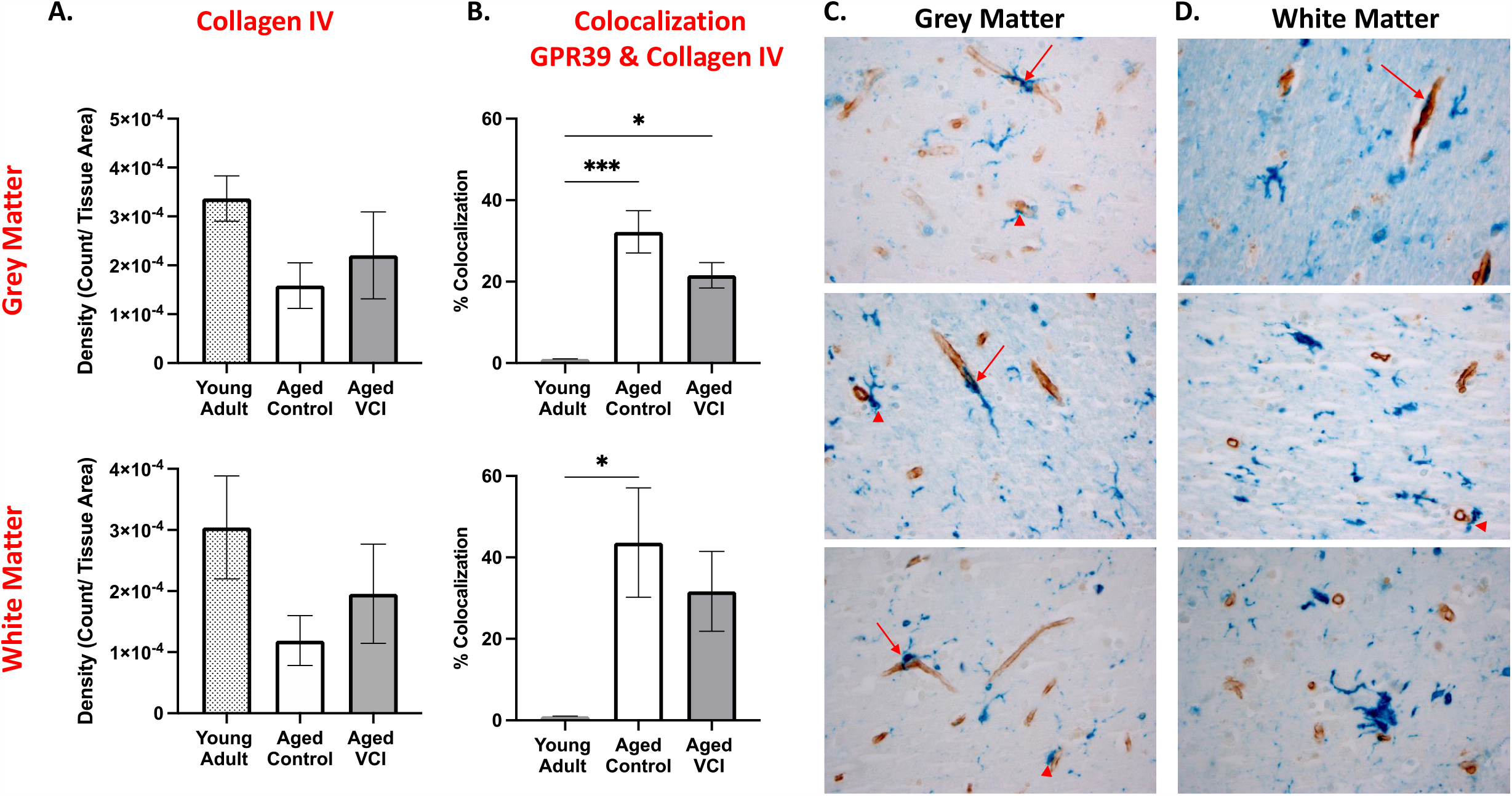
Peri-capillary localization of GPR39 in aged human dlPFC. **A**, Quantification of collagen IV expression density and **B**, colocalization of GPR39 with collagen IV, in both grey (top) and white (bottom) matter of young and aged control and aged VCI human dlPFC. Representative images of collagen IV (brown) and GPR39 (blue) labeling in human dlPFC grey, **C**, and white matter, **D**, arrows indicate colocalization of GPR39 with collagen IV, arrowheads indicate close proximity of GPR39 with collagen IV. ^*^ p < 0.05, ^**^ p < 0.001, 1-way ANOVA, data represent mean ± SEM (n=5-15/group).

We then investigated the cellular identity of the second population of GPR39-positive cells within the parenchyma. Morphologically these cells resemble microglia. We therefore performed double-labeling for GPR39 and the microglia/ macrophage marker CD68 (Figure 3). The density of CD68-positive cells as well as cells positive for both GPR39 and CD68 was higher in white matter than grey matter (Figure 3 C-D). We found that CD68 density was unaltered by age or dementia status (Figure 3A-B). While there were no significant differences in density of CD68-GPR39 co-localization between groups in grey matter, we observed a significant increase in the co-localization density in the aged VCI white matter compared to both young adult and age-matched control, increasing from 8.633 ×10^−6^ ± 4.68 ×10^−5^ in young adult and 2.49 ×10^−5^ ± 0.57 ×10^−5^ in aged control to 6.16 ×10^−5^ ± 1.34 ×10^−5^ count/tissue area in aged VCI dlPFC (P<0.05, 1-way ANOVA with Tukey’s multiple comparisons test; Figure 3B). Overall, we observed high levels of colocalization of GPR39 with CD68, with 60-85% of CD68-positive area also positive for GPR39 in both aged groups (Figure S1). Finally, we performed double immunolabeling of GPR39 with NeuN (Figure S2), smooth muscle actin (SMA) and glial fibrillary acidic protein (GFAP), where no co-localization was observed (not shown), indicating that GPR39 is not expressed by neurons, smooth muscle cells or astrocytes, respectively, in the human dlPFC.

**Figure 3.**
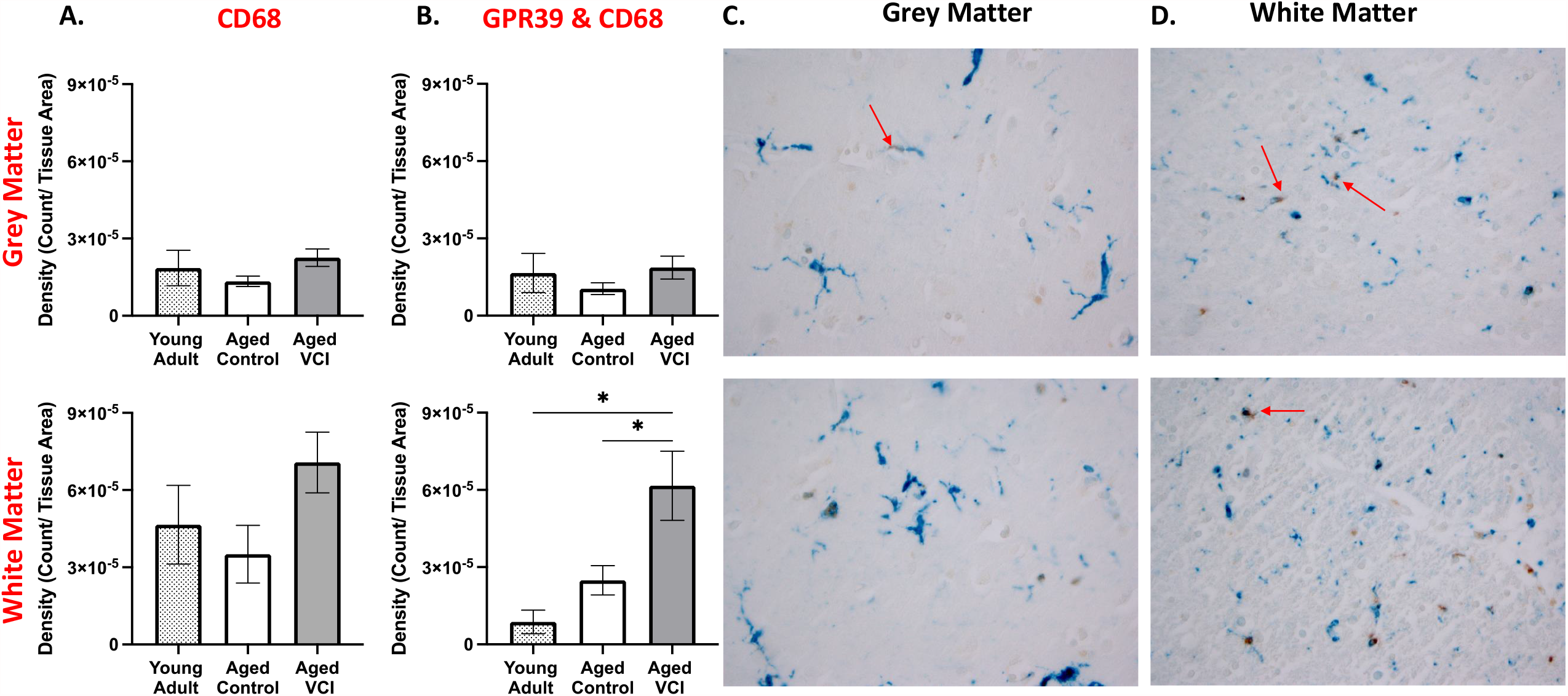
Microglial localization of GPR39 in aged human dlPFC. **A**, Quantification of CD68 expression density and **B**, density of colocalized GPR39 with CD68, in both grey (top) and white (bottom) matter of young and aged control and aged VCI human dlPFC. Representative images of CD68 (brown) and GPR39 (blue) labeling in human grey, **C**, and white matter, **D**, arrows indicate colocalization of GPR39 with CD68. ^*^p < 0.05, 1-way ANOVA, data represent mean ± SEM (n=5-15/group).

### GPR39 polymorphisms in control and MCI subjects and correlation with WMH burden

We next sought to determine whether polymorphisms in the gene encoding for GPR39 are associated with MCI diagnosis. We determined the presence of two single nucleotide polymorphisms (SNPs) [15] and wild type GPR39 in MCI and control subjects. In agreement with previous studies [15],we observed that the two SNPs were in high linkage disequilibrium with each other (SNPs were separated by a small distance genetically), allowing us to group genotypes into WT, homozygous SNP (both SNPs are homozygous) or heterozygous SNP (both SNPs are heterozygous). We found that in our cohort, there was no statistically significant difference between control and MCI groups in distribution of genotypes (Fisher’s exact test, p = 0.102). Importantly, however, out of a total of 78 subjects, only 5 carried the homozygous SNP; these were exclusively in the MCI group (Figure 4A).

**Figure 4.**
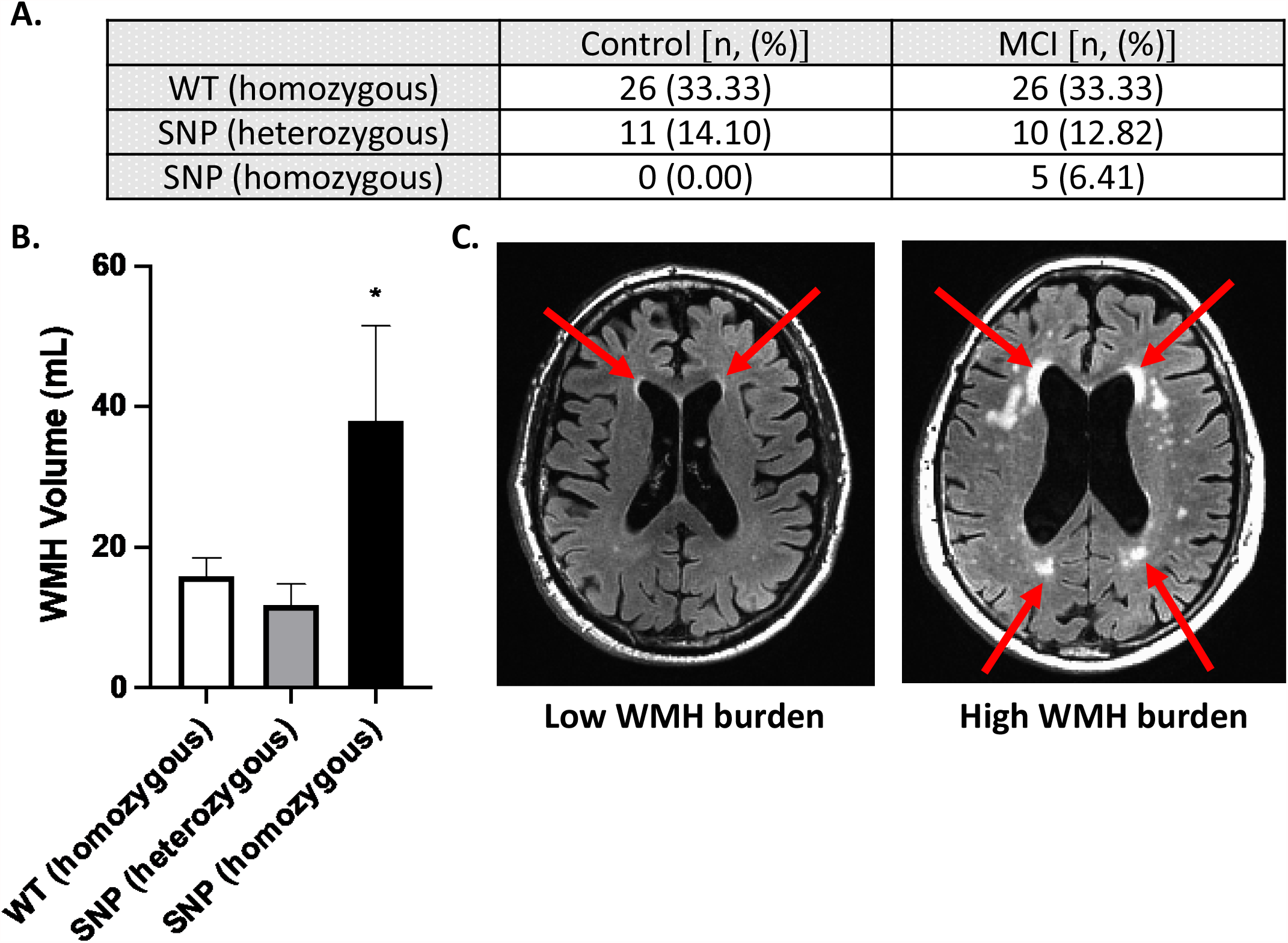
GPR39 single nucleotide polymorphisms in control and VCI subjects. **A**, Number of subjects carrying one of two GPR39 polymorphisms, or the WT GPR39 gene, in control and VCI groups. Fisher’s exact test determined lack of significance in genotypes between control and VCI subjects (p = 0.102). **B**, Quantification of WMH volumes by GPR39 genotype, regardless of clinical status. Significance was determined by one-way ANOVA with Tukey’s multiple comparisons test, ^*^ indicates p < 0.05 compared to both other groups. **C**, Representative 3T MRI scans of subjects with low (left) and high (right) WMH burden. Arrows indicate areas of WMH.

We then determined if GPR39 SNPs are specifically linked to WMH burden, which we previously demonstrated to be higher in VCI [7]. We found that carriers of the homozygous SNPs, which are only present in the MCI group, had significantly higher WMH volume compared to either WT or heterozygous SNP carriers (Figure 4B). WMH volume was 15.87 ± 2.59 mL and 11.90 ± 2.91 mL in WT and heterozygous SNP carriers, respectively, compared to 38.03 ± 13.46 mL in the homozygous SNP carriers (P<0.05, one-way ANOVA with Tukey’s multiple comparisons test).

## Discussion

This is the first study of GPR39 in human brain, and the first to implicate GPR39 in vascular cognitive impairment. We made three novel observations: 1) GPR39 is abundantly expressed throughout the human dlPFC; 2) GPR39 is expressed in two cell populations in the aged human brain; CD68-positive microglia/macrophages and CD68-negative cells in close proximity to capillaries, likely pericytes; 3) Homozygous GPR39 SNPs correlate with WMH burden and found exclusively in our study in individuals with cognitive impairment. The findings implicate GPR39 SNPs as a risk factor for VCI, and suggest that GPR39 may play a role in the pathogenesis of VCI and serve as a therapeutic target.

GPR39 is an orphan G-protein coupled receptor that belongs to the ghrelin receptor subfamily, which in addition to ghrelin includes motilin, neurotensin, neuromedin U (NMU) receptors [16]. The natural ligand for GPR39 remains unknown. Obestatin, a ghrelin-associated peptide derived from posttranslational processing of the prepro-ghrelin gene, was once proposed to be the GPR39 endogenous agonist, but this conclusion has been refuted[17]. Like ghrelin and NMU receptors, GPR39 is constitutively active[18]. Zinc ion (Zn^2+^) has also been shown to activate GPR39 receptor[19], and to modulate receptor activation [20]. GPR39 has been implicated in a variety of functions and disease conditions[21]. Because GPR39 is in the same family as ghrelin (the “hunger hormone”), studies initially investigated its role in insulin secretion, type 2 diabetes, food intake and weight control[21-24]. More recent studies have investigated its role in a range of brain disorders, including epilepsy[25], alcohol use disorders[26], anxiety and depression[27], and protection from neuronal cell death[28]. Of interest to the current study, GPR39 has been implicated in cognitive impairment associated with AD[9].

Regional expression of GPR39 transcript and protein have been reported in both mouse and human brain[16], with expression detected in multiple brain regions, including the amygdala, hippocampus and frontal cortex [29-31] [32]. Most of these studies, however, have been conducted using Northern or Western blot analysis of tissue homogenates, which precluded the identification of the specific cell type expressing GPR39. The few studies that used in-situ hybridization or immunohistochemistry to localize GPR39 in mouse brain have reported expression in neurons[33] [34]. Our study in human dlPFC does not find neuronal expression, but rather expression in microglia and peri-capillary cells. Furthermore, we find that perivascular expression is increased in both the aged and aged MCI dlPFC, compared to young, whereas microglial expression is increased in MCI dlPFC compared to both aged and young control.

A recent study demonstrated that CD68 immunoreactivity is increased in human brains corresponding with higher level of white matter lesions [35]. Based on this and other observations, Waller et al. concluded that microglial activation is a prominent feature of age-associated white matter lesions [35]. As the resident brain immune cell, microglia may be increased in number or activated by oxidative stress associated with SVD, which contributes to the pathogenesis of dementia [36]. Our finding that a high percentage of microglia in the aging brain express GPR39, and that the density of GPR39-positive cells is increased in the white matter of MCI dlPFC implicates GPR39 in the neuroinflammation underlying aging-related cognitive decline. In a model of chronic cerebral hypoperfusion, a zinc complex has been shown to ameliorate cognitive impairments and reduce microglial activation [37], raising the possibility that this effect may be mediated by GPR39. Although the GPR39-expressing CD68 cells in our study appear morphologically microglial, the possibility that these cells are blood derived macrophages cannot be discounted. CD68 labels both microglia and macrophages, although combinations of CD68 with other markers attempt to distinguish the two cell types, a definitive identification is difficult, particularly since phenotype of these cells is age-and function-specific[38]. For example, TMEM119 may be used in combination with CD68, however its expression is not stable throughout the lifespan or activation state of the microglia[39], while the traditional microglial marker, Iba1 does not appear to be expressed in all microglial populations [35]. Combinations of markers and their expression levels (TMEM119, CD45, Iba1, CD68) may help distinguish between the cell types [39], such quantitative analysis of immunolabeling intensity, with multiple markers is challenging on human tissue.

Perivascular expression of GPR39 in our study suggest a role in pericyte function. Brain peri-capillary pericytes, lodged between capillary endothelium and astrocytic end feet, form part of the neurovascular unit. Analogous to smooth muscle cells in arteries and arterioles, they are contractile and control blood flow at the capillary level [40-43]. It is therefore unsurprising that experimentally, loss of pericytes reduces brain capillary perfusion, leading to chronic perfusion stress, hypoxia, and progressive neurodegeneration, including white matter abnormalities suggestive of SVD [44] [45]. In humans, alterations in pericyte function contribute to impaired neurovascular coupling with aging, believed to contribute to SVD and subsequent cognitive decline[46-48]. We demonstrate a robust increase in perivascular expression of GPR39 in aged human dlPFC, both control and MCI, compared to young adult, in cells morphologically consistent with pericytes, with expression located around small, but not large vessels. We also observe a trend towards decreased capillary density in the aged dlPFC, in both control and MCI brain, consistent with previous findings in other brain regions [49]. Although we did not observe differences in expression levels between MCI and aged-matched control subjects, this does not discount that receptor function may be altered between the two groups. The function of GPR39 in these cells, and whether it may contribute to age-related vascular alterations, remains to be determined.

Our study demonstrates for the first time that homozygous SNPs in the gene encoding for GPR39 are associated with increased WMH burden. We have previously demonstrated that acceleration of WMH burden, a common indicator of cerebral SVD in the elderly, occurs prior to symptomatic MCI [6]. Although genotyping for the SNPs did not distinguish between MCI and control groups, subjects homozygous for both SNPs fell exclusively into the MCI group; it is possible that with increased sample size (there were only 5 homozygous out of 78 subjects), we may observe a correlation between these SNPs and the clinical diagnosis. In our cohort we observed linkage disequilibrium in every subject, an observation also made by Slavin et al.[15], indicating that these two particular SNPs are close to each other. If validated in a larger study, these SNPs can therefore be used as biomarkers to identify individuals at higher risk of developing WMH in later life and adopting disease-modifying strategies prior to WMH acceleration and dementia onset. Although GPR39 SNPs have not been implicated in cognitive impairment, previous studies implicated variants in GPR39 in multiple conditions including hypertension, nerve injury[50] and pediatric obesity[51].

In summary, we report that GPR39 is expressed in the dlPFC of the aged human brain in microglia/macrophages and peri-capillary cells resembling pericytes: the colocalization of GPR39 with capillaries is increased in the aged dlPFC compared to young adult, and the density of GPR39-positive microglia is higher in dlPFC white matter in VCI vs. age-matched and young adult control brain. We also demonstrate that homozygous GPR39 SNPs are associated with increased WMH volume. These findings suggest that GPR39 may play a role in regulating inflammation and capillary function in the aging brain, and also identify homozygous SNPs as a potential biomarker for development of SVD and MCI.

## Acknowledgements/ Conflicts/ Funding

We would like to acknowledge expert technical assistance by staff in the OHSU Advanced Light Microscopy Core, supported by grant P30NS061800 and OHSU Biostatistics & Design Program (partially supported by the Oregon Clinical and Translational Research Institute [UL1TR002369]) for data analysis expertise. DNA quality assessment and qPCR assays were performed in the OHSU Gene Profiling Shared Resource, which receives support from the OHSU Knight Cancer Institute NCI Cancer Center Support Grant P30CA069533. We thank the Oregon Alzheimer’s Disease Research Center (NIA P30AG066518, P30AG008017) and the Department of Veterans Affairs.

NJA is co-inventor of GPR39-targeting compounds that have been licensed to a company with a commercial interest in the results; in the past 36 months, NJA co-founded, then sold shares in a company (Vasocardea), which was formed to develop GPR39-targeting drugs and diagnostic probes. He is also founder of another company (NeuvaRx), formed to develop GPR39-targeting drugs and diagnostic probes for CNS disorders. This potential conflict of interest has been reviewed and managed by OHSU. NJA has received royalties for a co-invention ([18F]FNDP for PET Imaging of Soluble Epoxide Hydrolase), licensed to Precision Molecular. NJA has also received compensation from NIH for grant reviews, and honoraria from academic universities for invited lectures. NJA has one patent issued (WO2017192854A1: Radiofluorinated FNDP for PET imaging of soluble epoxide hydrolase (sEH), and two other provisional patents are pending (Use of GPR39 probes and ligands for the diagnosis and treatment of cardiovascular disease, and biomarkers for detection of coronary artery disease and its management). KG has received consultant fees from OHSU for coding tools. Other authors have declared that no conflict of interest exists.

In the past 36 months the following authors have received support outside this project: CMD, Oregon Partnership for Alzheimer’s Research; JWN, NIH 1K01DK121737; LCS, NIH/NIA R01 AG056712; NJA, NINDS R01 NS108501.

This work was supported by NIH grant to NJA 1RF1AG058273-01.

