## Supplemental Figure 1 for "GPR39 Localization in Aging Human Brain and Correlation of Expression and Polymorphism with Vascular Cognitive Impairment"

**Fig S2. Double-labelling of GPR39 and neurons.**

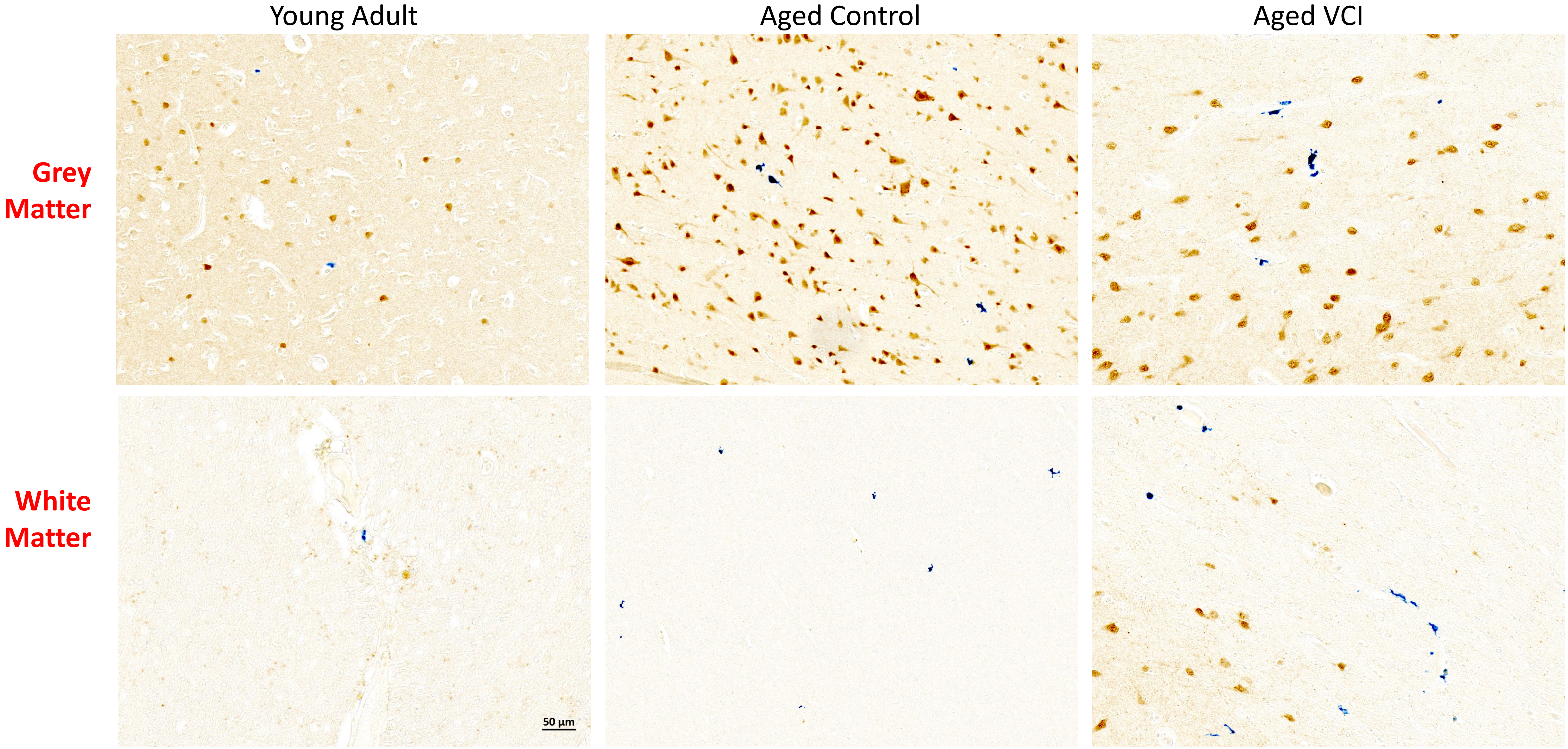


**Figure S1: Microglial colocalization of GPR39.** Quantification of colocalization of GPR39 with CD68, in both grey and white matter of aged control and VCI human brains. Data represent mean [illegible] SEM (n=13-15/group). Representative images shown in Figure 3.
