## Supplemental Figure 2 for "GPR39 Localization in Aging Human Brain and Correlation of Expression and Polymorphism with Vascular Cognitive Impairment"

**Figure S2: Double-labelling of GPR39 and neurons.**

Representative images of NeuN (brown) and GPR39 (blue) in human dlPFC grey and white matter.

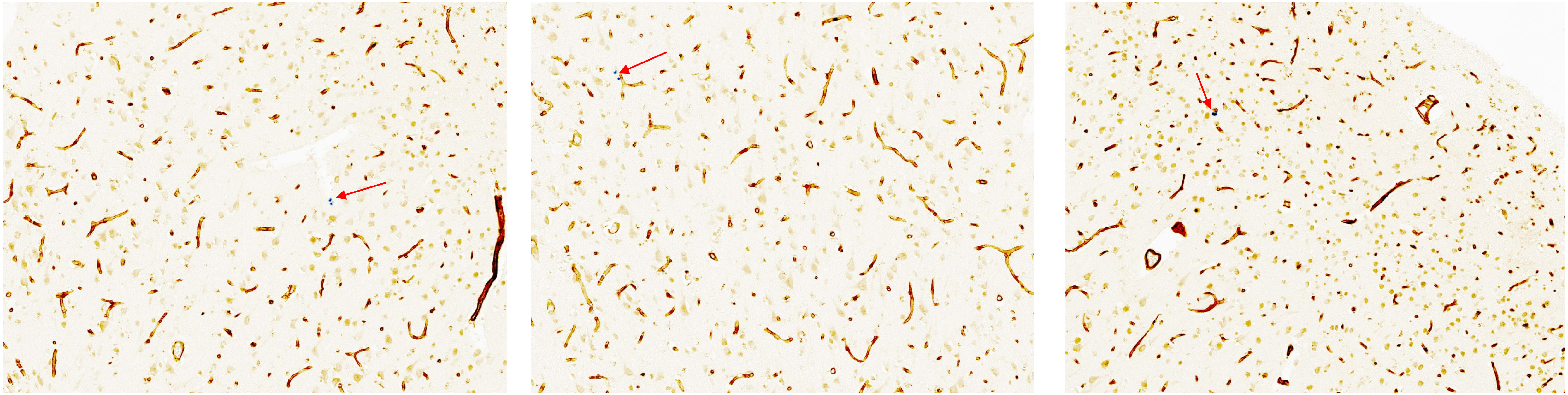
